# Engineering and evaluation of *Sinorhizobium meliloti* nodulation (*nod*) gene reporter systems in rhizobia and non-rhizobia

**DOI:** 10.1101/2025.11.03.686230

**Authors:** Chinh X. Luu, Barney A. Geddes

**Affiliations:** Department of Microbiological Sciences, North Dakota State University, Fargo, ND, USA

**Keywords:** *Sinorhizobium meliloti*, nodulation gene reporters, *Sinorhizobium-Medicago* symbiosis, symbiotic signal biosensors, flavonoid-responsive reporter, betaine-responsive reporter, non-rhizobia diazotrophs

## Abstract

Developing N□-fixing partnerships between diazotrophic microbes and non-legumes can enhance soil fertility and reduce dependence on synthetic fertilizers. Unlike legumes, non-legumes lack the genetic ability to form root nodule symbiosis with rhizobia but can form facultative associations with free-living diazotrophs. Engineering root nodule symbiosis in non-legumes remains a central aim in synthetic biology to enhance biological nitrogen fixation in cereals. Such a symbiosis will require specific symbiotic signaling that mimics the rhizobium-legume interaction. However, lack of effective tools for identifying compatible and engineerable microbial partners is a key challenge. To address this, we have developed inducible nodulation (*nod*) gene reporters to screen both rhizobial and non-rhizobial strains capable of expressing *Sinorhizobium meliloti nod* genes, which encode bacterial signals initiating nodule formation in legumes. The reporters include a superfolder GFP reporter controlled by the inducible *nod* box promoter (P*nodA*), plant signal-dependent activators *nodD1* and *nodD2*, and a constitutively *mScarlet-I* marker, named *nodD1*-P*nodA* and *nodD2*-P*nodA*. Their functionality was validated in various *S. meliloti* backgrounds using *in vitro* induction and two *in planta* induction approaches. These advancements facilitated the identification of both rhizobia and non-rhizobia capable of expressing *S. meliloti nod* genes, thereby supporting the development of synthetic N□-fixing symbioses in cereals.

## 1 INTRODUCTION

Integration of symbiotic and associative bacterial nitrogen fixation into crops represents a promising strategy for sustainable nitrogen management in agriculture (Haskett et al. 2022). Biological nitrogen fixation (BNF), in which N□-fixing microbes convert atmospheric nitrogen (N□) into bioavailable ammonia (NH□) under ambient conditions, is a natural process that offers a renewable and eco-friendly solution to reduce the reliance on synthetic nitrogen fertilizers in agriculture. BNF occurs in three ecological forms: symbiotic, associative, and free-living (Guo et al. 2023).

Among these, symbiotic nitrogen fixation (SNF) in legumes is particularly well-characterized and involves a highly coordinated interaction between host plants and rhizobia (Oldroyd 2013). This symbiotic process begins with a coordinated exchange of signals and nutrients between plants and bacteria. Legume roots secrete signaling compounds such as flavonoids, betaines, and other secondary metabolites into the rhizosphere. These signals activate rhizobial NodD proteins, which induce the transcription of nodulation (*nod*) genes and the production of Nod factors (NFs). Upon recognition of NFs, the plant initiates the formation of infection threads (ITs) and the development of root nodules. Rhizobia enter the root cells through ITs and differentiate into nitrogen-fixing bacteroids within nodules (Oldroyd 2013). The plant supplies carbon sources to the bacteroids, which in turn reduce atmospheric nitrogen into ammonia (Udvardi and Poole 2013). The *Sinorhizobium*-*Medicago* symbiosis serves as a model system for studying SNF (Jones et al. 2007). In this interaction, *Sinorhizobium meliloti* responds to flavonoid and betaine signals from host plants, such as *Medicago sativa* (alfalfa) and *Medicago truncatula* (barrelclover), by activating NodD1 and NodD2, respectively. These transcriptional regulators bind to a specific sequence upstream of the nodulation (*nod*) gene operons to initiate NF production for colonization in legumes (Barnett and Long 2015). *S. meliloti* also possesses a third NodD protein, NodD3 which activates the expression of nodulation genes and the subsequent synthesis of NF in a flavonoid-independent manner (Barnett and Long 2015).

In contrast to legumes, cereal crops such as maize, wheat, and rice lack the genetic machinery to form root nodules and establish intimate symbiotic relationships with diazotrophic bacteria. Current strategies to enhance nitrogen delivery in cereals focus on promoting associative interactions or transferring SNF pathways into non-leguminous hosts (Guo et al., 2023). Though many cereal-colonizing microbes are capable of fixing nitrogen in association with cereals (Mus et al. 2016; Haskett et al. 2022), these taxa do not possess the symbiotic signaling machinery required to initiate nodulation (NF production). Thus, engineering an orthogonal symbiotic signaling in cereal-colonizing microbes through NF-dependent (Geddes et al. 2019; Haskett et al. 2022) or independent (Haskett et al. 2025) cascades will be a critical contribution to synthetic root nodule symbiosis in cereals. A major challenge in these approaches is the identification of compatible and engineerable microbial partners.

To tackle this problem in the context of engineering orthologous NF-based signaling, we developed and validated inducible biosensors with two aims. First, to provide insight into the activation of symbiotic signaling in the native strain *S. meliloti*, and second to rapidly identify rhizobial and non-rhizobial strains capable of expressing *S. meliloti* nodulation genes and producing *S. meliloti* NFs in response to plant-derived symbiotic signals. The *nodD1*-P*nodA* system was used as a flavonoid-responsive biosensor, while *nodD2*-P*nodA* was served as a betaine-responsive reporter. Using the symbiotic signal biosensor, we provided key insights into how flavonoid-mediated NodD1 and betaine-mediated NodD2 regulate the *S. meliloti* NF production during the early stages of *Sinorhizobium-Medicago* symbiosis. The independent expression of the flavonoid biosensor made it suitable for the identification of *S. meliloti nod* gene expression capability in rhizobia and non-rhizobia. Incorporating these plant signal biosensors into high-throughput screening workflows for diazotrophs capable of expressing synthetic N fixation genes is expected to facilitate the integration of biological nitrogen fixation (BNF) capacity into non-leguminous crops, thereby enhancing crop productivity and improving soil health.

## 2 MATERIALS AND METHODS

### 2.1 Bacterial strains and culture conditions

All bacterial strains were listed in Table 1. *Escherichia coli* strains were grown on solid and liquid Luria-Bertani (LB) medium at 37 °C. For *E. coli* ST18 strain culture, 50 µg·mL^-1^ of 5-Aminolevulinic acid Hydrochloride (ALA) was supplemented on the LB medium. *S. meliloti* strains were cultured on solid and liquid LB medium with the supplement of 2.5 mM of MgSO_4_ and CaCl_2_ (LBmc) at 30 °C. Rhizobia strains were grown on solid and liquid Tryptone-Yeast extract (TY) medium supplemented with CaCl_2_ and biotin at 28 °C. Tryptic Soy Agar (TSA) was used for culturing non-rhizobia strains at 28 °C. The bacterial strains were cultured and stored at −80 °C in the medium solution containing 8% Dimethyl sulfoxide (DMSO) until use.

**Table 1.**
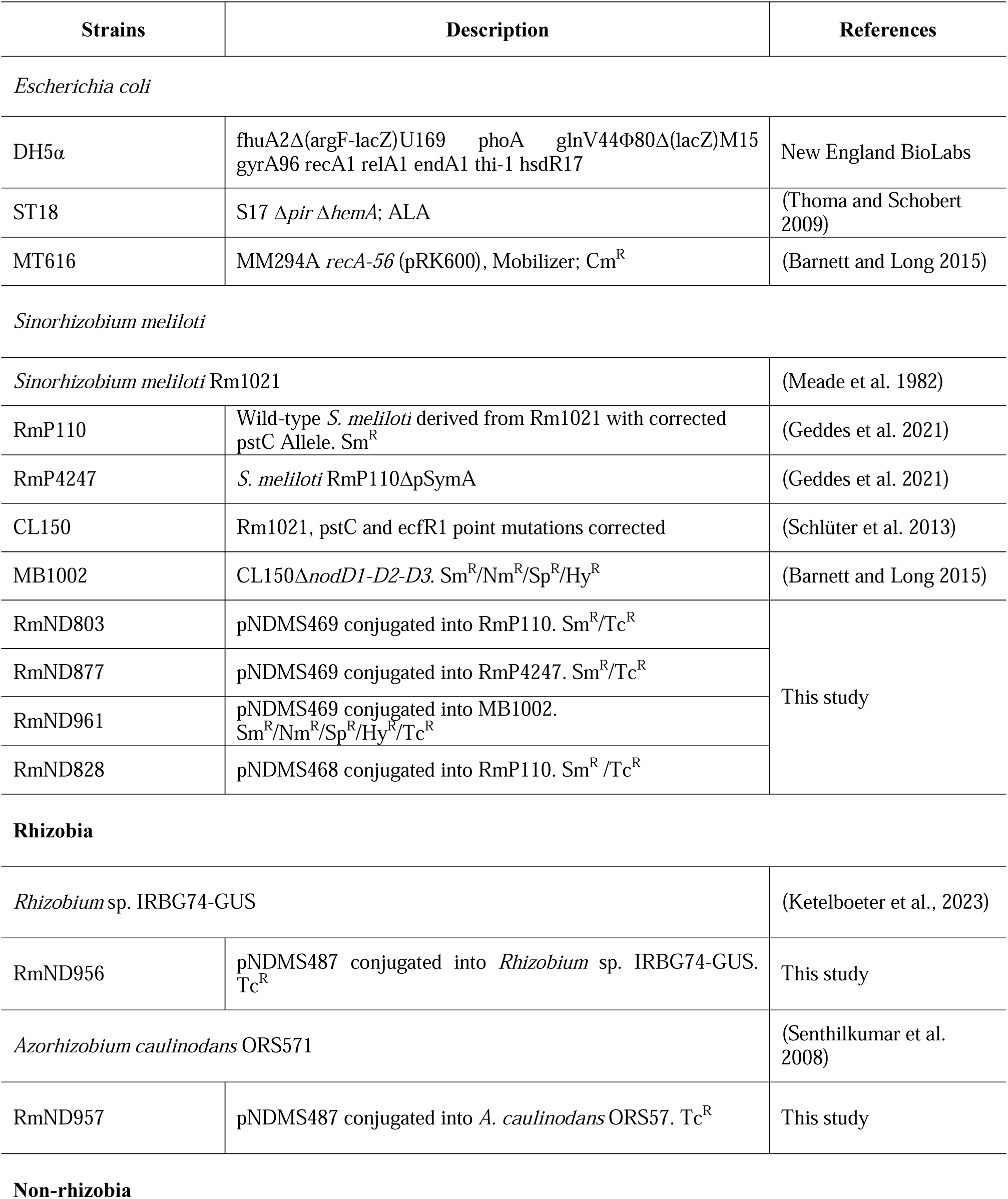
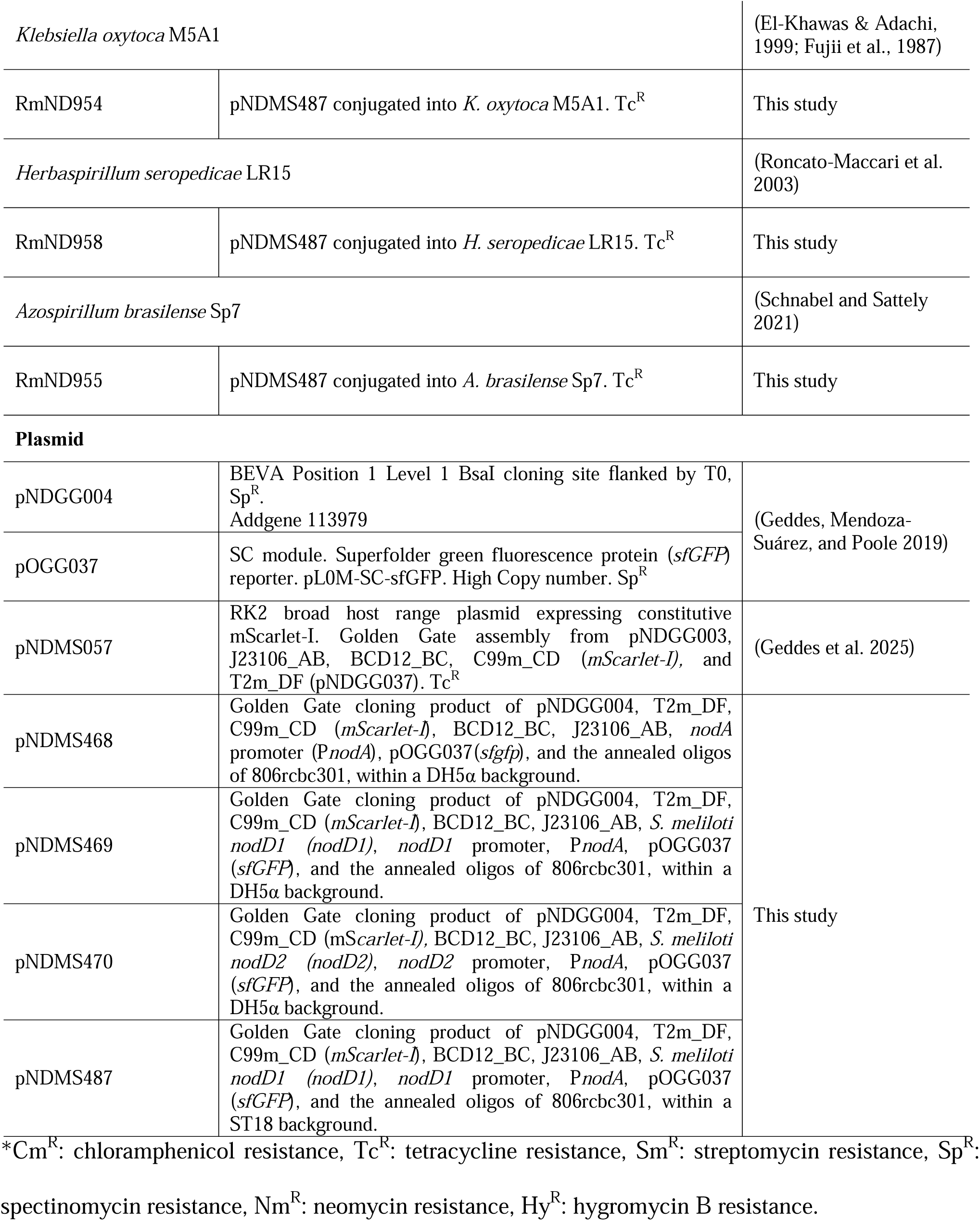
Bacterial strains and plasmid used in the study.

The following concentrations of antibiotics were used 2.5 µg·mL^-1^ chloramphenicol (Cm), 5 µg·mL^-1^ tetracycline (Tc), 200 µg·mL^-1^ streptomycin (Sm), 200 µg·mL^-1^ neomycin (Nm), 5 µg·mL^-1^ spectinomycin (Sp) and 5 µg·mL^-1^ hygromycin B (Hy).

### 2.2 Construction of *S. meliloti nod* reporters

To construct pNDMS469 (*nodD1-*P*nodA*), *mScarlet-I* under the control of a constitutive promoter (P_J23106_) was amplified from pNDMS057 (Geddes et al. 2025), and combined with *S. meliloti nodD1* (*nodD1*) including 250-bp upstreams of *nodD1* that contained the *nodA* promoter (P*nodA*), and superfolder green fluorescent protein (*sfGFP*) gene downstream of *PnodA*, terminated by a single unique Barcode-ID (806rcbc301) (Schumacher et al. 2025) (**Figure 1A, left**). All PCR primers are listed in Supplementary Table S1.The parts were combined via amplification with compatible BsaI cutsites that facilitated linear assembly in the order described above. Alternatively, for the construction of pNDMS470 (*nodD2-*P*nodA*), *mScarlet-I* driven by P_J23106_ was combined with *S. meliloti nodD2* and its 1120-bp upstream region (P*nodD2*), along with the P*nodA*-sfGFP fusion. In contrast, pNDMS468 (P*nodA*) included *mScarlet-I* under the P_J23106_ promoter and the P*nodA*-sfGFP fusion only. NEB Golden Gate Assembly Kits (BsaI-HF-v2) were used as a pre-mixed reaction mix with the DNA parts and ddH_2_O. The Golden Gate cloning reactions were performed in a thermocycler as follows: 30 cycles of 37 °C for 1 min and 16 °C for 1 min, with heat inactivation at 60 °C for 5 min. The resulting reaction mixes were stored at −20 °C before thawing and using to transform chemically competent *E. coli* DH5α. Successful transformants were selected using LB^Tc^ plates. Plasmids were verified by diagnostic restriction digest and for long read Oxford Nanopore sequencing by Plasmidsaurus to confirm that no errors were present. The full sequence of each plasmid is available in Supplementary Table S2. The plasmids were then introduced to a conjugation donor strain ST18 by transformation.

**Figure 1.**
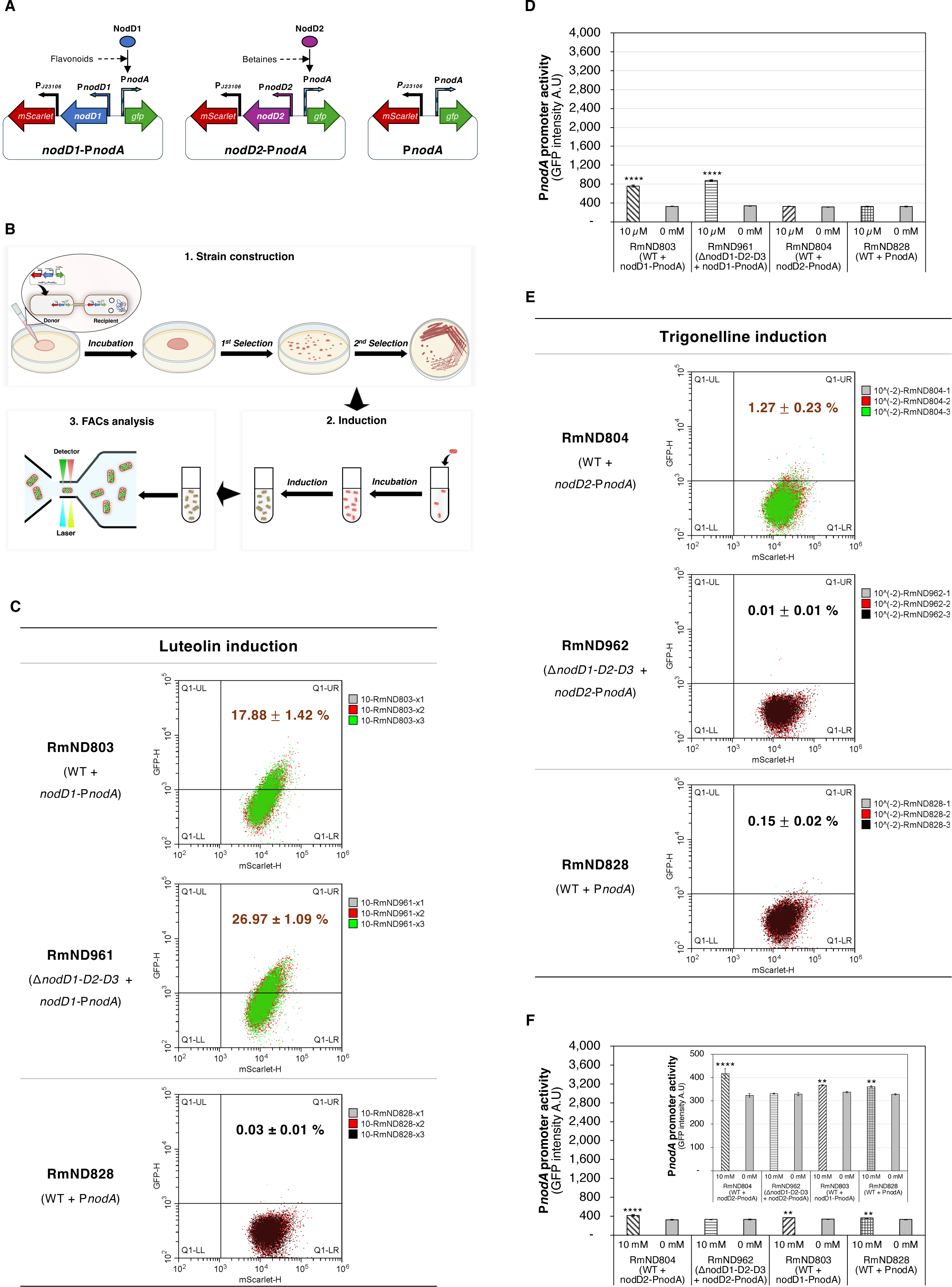
Design and validation of symbiotic signal-dependent *S. meliloti nod* reporters. (A) Schematic designs of biosensor constructs: flavonoid-responsive *nodD1*-P*nodA*, betaine-responsive *nodD2*-P*nodA*, and P*nodA* reporter constructs. (B) Workflow for strain construction, *in vivo* induction, and flow cytometry (FACs) analysis. (C) sfGFP and mScarlet-I fluorescence intensity analysis for populations of *S. memiloti* RmND803 (wild-type with the *nodD1*-P*nodA*) and RmND961 (Δ*nodD1-D2-D3* mutant with *nodD1*-P*nodA*), compared to the control strain RmND828 (WT with P*nodA*), after 4 h induction with 10 µM luteolin. A fluorescence-based gate was established using the control strain to identify sfGFP-positive cells. Data represent the average proportion of sfGFP-bacterial population (mean ± standard deviation) from three independent experiments, with 10,000 mScarlet-I-expressing cells analyzed per experiment. (D) Quantification of flavonoid-dependent activation of the *nodD1*-P*nodA* reporter, shown as sfGFP fluorescence intensity from FACS analysis. RmND828 (P*nodA-*carrying strain) population induced by luteolin served as the negative control. Data represent the green fluorescent intensity (mean ± standard deviation) from three independent experiments. (E) FACs analysis of RmND804 (WT with *nodD2*-P*nodA* construct), RmND962 (mutant with *nodD2*-P*nodA*), and the control strain RmND828 under 10 mM trigonelline induction. The same gating strategy was applied to identify sfGFP-positive cells. Data represent the mean ± standard deviation of sfGFP-positive populations from three independent experiments, each analyzing 10,000 mScarlet-I-expressing cells. (F) Green fluorescence readout for betaine-dependent activation of *nodD2*-P*nodA* reporter based on the FACs analysis. RmND828 (P*nodA-*carrying strain) population induced by trigonelline served as the negative control. Data represent the green fluorescent intensity (mean ± standard deviation) from three independent experiments.

### 2.3 Conjugating the *nod* reporters into rhizobia and non-rhizobia

Conjugations between *E. coli* and *S. meliloti* were conducted through triparental mating using *E. coli* MT616 as a helper strain (Finan et al. 1986). Meanwhile, *E. coli* ST18 served as a donor strain in biparental mating with other recipient strains (Thoma and Schobert 2009). Briefly, overnight cultures of each strain were mixed with a 1:1:1 or 1:1 ratio and spotted onto culture plates (LBmc or TY or TSA plates) without antibiotics, then incubated for 24 h at 30 °C. The 24-h incubated mating spots were resuspended in 1 mL of physiological saline (0.9% NaCl), diluted serially (10^-1^, 10^-2^, and 10^-3^), and then plated onto selection plates with appropriate antibiotics. The plates were incubated at 30 °C until the appearance of visible bacterial colonies. Single colonies from selection plates were streaked to obtain purified single colonies.

### 2.4 *In vitro* induction of the *nod* reporters

A single colony was inoculated in 5 mL of media containing standard working concentrations of relevant antibiotics and cultured at 30 °C, 250 rpm for 24-72 h. After the cultivation, the optical density at 600 nm (OD_600_) was measured using a spectrophotometer. The bacterial suspension was diluted to a final OD_600_ of 0.2 in M9 minimal salt solution, which was supplemented with the appropriate antibiotics and 10 µM of luteolin for 4 h or 10 mM trigonelline for 6 h.

### 2.5 *In planta* induction of the *nod* reporters

Plants used in this experiment included *Medicago sativa* cv. Iroquois and *Lotus japonicus* cv. *gifu*. *M. sativa* seeds were surface sterilized with 95% ethanol for 5 min and with 2.5% NaClO for 20 min. After sterilization, the seeds were rinsed 10x with sterile water for one hour, with the water being replaced every 15 min. Lotus seeds were treated with 98% sulfuric acid for 12 minutes. The seeds were rinsed ten times with sterile water. Next, the seeds were soaked in 15 mL of a 2.5% bleach solution for 20 min, followed by an additional ten rinses with sterile water. Finally, the seeds were soaked in sterile water for 2 to 4 h to complete the sterilization process.

For the germinating seed induction, the seeds were then added in a bacterial suspension with an OD_600_ of 0.2 at room temperature for 1 h, before being placed onto 0.8 % agar plates. The seeds were grown in Conviron Gen1000 or Gen2000 growth chambers with LED lights for germination. The environmental settings in the chambers were programmed to simulate an 18-hour daytime period at 21 °C with full light intensity, followed by a 6-hour nighttime period at 17 °C in complete darkness.

For the root induction of bacteria carrying the bioreporters, a paper-based growth (Pa-Gr) system was employed (Le Marié et al. 2014). In brief, sterilized *M. sativa* and/or *L. japonicus* seeds were placed on 0.8% agar, covered with aluminum foil, and incubated in the dark for 3 d. The foil was then removed, and incubation continued for 3 d. For each Pa-Gr file setup, three sprouts were arranged between blue and brown paper sheets, and this arrangement was repeated to obtain 15 plants per file (5 plants × 3 rows). The papers were secured using binder clips and inserted into the sterilized plastic envelopes, followed by the addition of 800 mL of sterilized H O and 10 mL of 10X Jensen’s medium. Each row was inoculated with 5 mL of a cultured bacterial suspension with an OD_600_ of 0.2, resulting in a total of 15 mL per file. The files were then placed in the plant growth chambers.

After 1-14 dpi, 25 germinating seeds or 5 plants were collected and placed into a 50 mL tube containing the phosphate-buffered saline (PBS) buffer plus 0.02% Silwet. The tubes were rotated at 70 rpm at room temperature for 1 h before filtering via 100 µm strainers to collect bacterial suspension. Subsequently, the reporter expressions in bacteria were analyzed by a flow cytometry (FACs) platform.

### 2.6 Fluorescence measurements by flow cytometry (FACs) platform and microscopic observations

The fluorescence expression levels were measured by Beckman Cytoflex S flow cytometer (Beckman Coulter, Inc., Indianapolis, IN, USA). The yellow light detector with a band path of 585±42 nm was set to measure the mScarlet-I (569 nm excitation and 593 nm emission) intensity.

The blue light detector was set at 525±40 nm to detect superfolder sfGFP intensity (488 nm excitation and 510 nm emission). Before the FACs analysis, the bacterial solution was diluted in the PBS buffer to adjust the speed settings for an average of 2,000 events per second. The flow speed was set at the lowest speed of 10 µL·min^-1^. Gates for mScarlet-I-positive populations were set using *S. meliloti* RmP110 and *E. coli* DH5α as negative controls, while sfGFP-positive gates were established using *S. meliloti* RmND828 under non-inducing conditions. Data output was analyzed by CytExpert (Beckman Coulter, Inc., Indianapolis, IN, USA).

For microscopic observations, root samples were collected, placed on a slide and mounted in a 50% glycerol solution, which serves as a mounting medium for fluorescence microscopy. The samples were then visualized using a Mica Microhub system (Leica Microsystems, Inc., Deerfield, IL, USA). In confocal mode, laser excitation at 488 nm was used for sfGFP and 561 nm for mScarlet-I, with fluorescence signals detected by the HyD FS FluoSync™ detector. In widefield mode, LED excitation at 470 nm and 555 nm was applied, respectively, and fluorescence signals were captured using the CMOS FluoSync™ detector.

### 2.7 Statistical analysis

A one-way ANOVA test with Dunnett’s multiple comparison test was used to establish significant differences between replicates. All data shown was significant as determined by one-way ANOVAs. In the figures, results from post hoc significance testing are indicated as *P < 0.0332, **P < 0.0021, ***P < 0.0002, ****P < 0.0001.

### 2.8 Phylogenetic analysis

For building a phylogenetic tree of model rhizobia and non-rhizobia, 16S rRNA sequences were obtained from NCBI GenBank. Sequences were globally aligned, and pairwise identity was calculated using a binary scoring scheme (1.0 for matches, 0.0 for mismatches). A genetic distance matrix was generated using the Tamura-Nei model, and a phylogenetic tree was constructed with the Neighbor-Joining method.

## 3 RESULTS

### 3.1 Construction of *S. meliloti nod* reporters

Identifying suitable and engineerable microbes is a key task in the development of synthetic N□-fixing symbioses in legumes and non-legumes. With the goal of screening for cereal-colonizing, N□-fixing microbes that are compatible with NF-based signaling, we constructed plant signal-dependent biosensors based on the *Sinorhizobium-Medicago* symbiosis model. These biosensors consisted of a tunable P*nodA-sfGFP* reporter fusion activated by symbiotic signal-dependent transcriptional activators (the flavonoid-inducible NodD1 protein or the betaine-inducible NodD2 protein) and a constitutively expressed *mScarlet-I* fluorescent marker. The P*nodA-sfGFP* fusions were used for monitoring *S. meliloti* nodulation (*nod*) gene expression by sfGFP fluorescence, while the fluorescence of mScarlet-I served as a bacterial indicator. The resulting constructs were designated as *nodD1*-P*nodA* and *nodD2*-P*nodA* (**Figure 1A**). A control construct, referred to as the P*nodA*, included only the P*nodA-sfGFP* reporter and *mScarlet-I*, without any transcriptional activator (**Figure 1A right**).

These reporters were then introduced into various *S. meliloti* genetic backgrounds to validate their inducibility and functionality (**Figure 1B**). The *nodD1*-P*nodA* reporter was conjugated into *S. meliloti* RmP110 (wild-type, WT) and MB1002 (Δ*nodD1-D2-D3* mutant), resulting in the conjugated strains RmND803 and RmND961, respectively. Similarly, the *nodD2*-P*nodA* construct was mobilized into the same backgrounds, generating strains RmND804 and RmND962. As a control, RmND828, the WT strain carrying the P*nodA* reporter, was generated. These strains were subsequently subjected to both *in vitro* and *in planta* induction assays.

### 3.2 *In vitro* induction for the *nod* reporters

To initially validate the functionality and inducibility of the symbiotic signal-responsive biosensors, we performed *in vitro* induction assays using plant-derived signaling compounds. These signaling compounds stimulate the transcriptional regulators NodD1 or NodD2 to activate *S. meliloti nod* gene transcription (Peck, Fisher, and Long 2006; Phillips et al. 1992). Specifically, the flavonoid luteolin was employed to activate the flavonoid-dependent *nodD1*-P*nodA* biosensor, while the plant-derived alkaloid trigonelline was used to activate the *nodD2*-P*nodA* reporter system. The *S. meliloti* strains harboring the reporter construct were cultured to the early stationary phase and subsequently induced with one of the selected plant signal compounds. The P*nodA* reporter activity was assessed by measuring sfGFP expression in mScarlet-I-labeled strains using a flow cytometry (FACs) platform (**Figure 1B**). The intensity of sfGFP from FACs analysis served as a relative measure of P*nodA* promoter activity.

Luteolin induction resulted in tunable activation of the *nodD*1-P*nodA* biosensor in *S. meliloti* strains RmND803 (WT harboring *nodD1*-P*nodA*) and the *nodD*-lacking background RmND961 (Δ*nodD1-D2-D3* mutant carrying *nodD1*-P*nodA*) (**Figure 1C and Supplementary figure S1A**). After 4-h induction, approximately 17.88 ± 1.42 % of RmND803 cell population and 26.97 ± 1.09 % of RmND861 cells exhibited sfGFP intensities exceeding the 99th percentile of the control strain RmND828 (WT with P*nodA*). No activation was observed in the absence of luteolin and in RmND804 (*nodD2*-P*nodA* carrying strain) under the same induction condition (**Supplementary figure S1A**). Statistical analysis of replicated treatments based on the sfGFP intensity further confirmed a significant increase in P*nodA* activity in RmND803 and RmND961 compared to RmND828 following luteolin treatment (**Figure 1D**). The P*nodA* activities of RmND803 and RmND961 under the luteolin induction were approximately 2.5-fold higher than that of the control strain RmND828, indicating the importance of including the regulatory protein in a proportional copy number to its promoter target.

Similarly, *nodD2*-P*nodA* activation was observed in RmND804 (WT harboring *nodD2*-P*nodA*), but not in RmND962 (*nodDs* null mutant with *nodD2*-P*nodA*), following induction with 10 mM trigonelline (**Figure 1E and Supplementary figure S1B**). After 6-h induction, only 1.27 ± 0.23 % of RmND804 cell population exhibited sfGFP fluorescence above the 99th percentile of the control strain RmND828. The reporter assays indicated that P*nodA* activity in RmND804 was 1.2-fold higher than that in the control strain (**Figure 1F**). No activation was detected in RmND962 and RmND803 (*nodD1*-P*nodA* carrying strain) under the same conditions.

Taken together, these findings validate the functionality and specificity of our inducible biosensors in response to flavonoid- and betaine-derived plant signals, though they point towards a much more robust activation of the *nodD1*-based sensor than the *nodD2*-based sensor with the inducers tested *in vitro*.

### 3.3 Germinating seed induction for the *nod* reporters

Following the successful *in vitro* validation of the symbiotic reporter systems, we further investigated whether the biosensors could report symbiotic responses during the early stages of microbe–plant interaction. To do this, we devised and employed a germinating seed induction approach to monitor biosensor activation under *in planta* conditions (**Figure 2A**). Germinating *M. sativa* seeds were treated with strains harboring the *nod* reporters and co-incubated on water-agar plates. After 1-, 2-, and 3-days post-inoculation (dpi), bacteria were detached from the root surface using PBS buffer solution supplemented with 0.02% Silwet. Reporter activation was assessed by detecting sfGFP expression in mScarlet-I-expressing strains via the FACs analysis.

**Figure 2.**
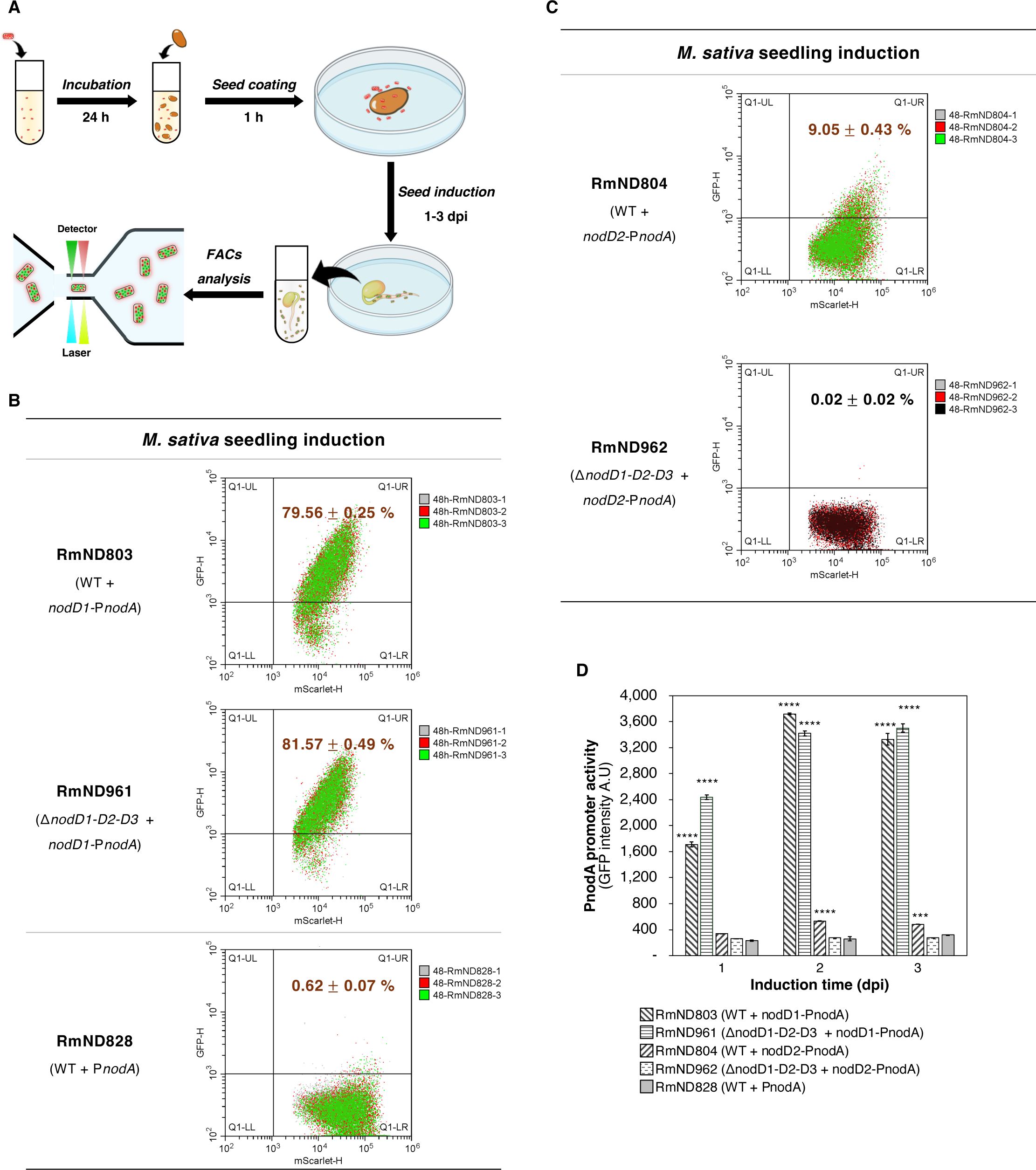
Germinating seed induction method for engineered *S. meliloti* strains carrying the *nod* reporters. (A) Experimental workflow for the germinating seed induction approach and subsequent FACs analysis of inoculated bacteria. (B-C) FACs analysis of engineered strains harboring the *nod* reporters after 2 days of the *M. sativa* seedling induction. The gating strategy established during in vitro experiments was applied to assess biosensor activation. Data represent the mean ± standard deviation of sfGFP-expressing cells from three independent experiments, each analyzing 10,000 mScarlet-I-expressing bacteria. (D) Activation of *nodD1*-P*nodA* biosensor in different *S. meliloti* backgrounds over a 3-day induction period. The control strain RmND828 (WT with P*nodA*) was included for comparison. Data represent the mean ± standard deviation of the green fluorescent intensity from FACS analysis across three independent experiments.

Our results revealed that P*nodA* activity in response to germinating seed exudates was significantly higher than that observed with single inducers (**Figures 2B-D and 1C-F**). For the activation of the flavonoid-dependent biosensor *nodD1*-P*nodA*, both RmND803 (WT carrying *nodD1*-P*nodA*) and RmND961 (Δ*nodD1-nodD2-nodD3* mutant carrying *nodD1*-P*nodA*) exhibited robust NodD1-mediated P*nodA* activity in response to natural alfalfa-derived signals throughout the 3-day induction period (**Supplementary Figure S2**). By day 2, 79.56 ± 0.25 % of RmND803 cells and 81.57 ± 0.49 % of RmND961 cells exhibited high sfGFP fluorescence, compared to a mere 0.62 ± 0.07 % of the control strain RmND828 (**Figure 2B**). By day 3, over 76% of RmND803 and RmND961 cells exhibited high sfGFP fluorescence, while the control strain exhibited fluorescence in approximately 3% of the population (**Supplementary Figure S2**). The biosensor quantification assays indicated that P*nodA* activities of RmND803 and RmND961 under the germinating seed induction were approximately 14-fold higher than that of the control strain RmND828 (**Figures 2D**). These activities were significantly greater than those induced under *in vitro* conditions (**Figure 1D**). For the activation of the betaine-dependent biosensor *nodD2*-P*nodA*, 9.05 ± 0.43 % of the RmND804 (WT with *nodD2*-P*nodA*) population showed high sfGFP fluorescence after the 2-d root induction. In contrast, RmND962 (Δ*nodD1-nodD2-nodD3* mutant with *nodD2*-P*nodA*) showed no detectable sfGFP expression (**Figure 2C and Supplementary Figure S2**). The reporter assays showed that P*nodA* activities of RmND804 under the germinating seed induction were up to 2-fold higher than that of the control strain, while no P*nodA* activation could be observed in RmND962 under the same conditions (**Figures 2D and Supplementary Figure S2**). Thus, the germinating seed induction approach effectively activated the *nodD1*-P*nodA* biosensor in *S. meliloti* backgrounds during the 3-day interaction period between engineered microbes and germinating *M. sativa* seeds, with induction showing a much greater magnitude of increase compared to induction with purified single inducers alone (**Figure 1F, 2D**). In contrast, the only induction observed by the *nodD2-PnodA* sensor in this system was dependent on the presence of other *nodD* proteins in the background genome.

### 3.4 Root induction for the symbiotic signal dependent *nod* reporter

In addition to germinating seed exudates, root exudates act as inducing signals for NF production, triggering infection thread and nodule formation in *Sinorhizobium–Medicago* symbiosis initiation (Peck, Fisher, and Long 2006; Le Marié et al. 2014). However, the dynamic nature of root exudation throughout plant development presents challenges for their collection and subsequent analysis of microbe–plant interactions. To overcome this limitation, a paper-based growth (Pa-Gr) system was employed for the first time to study rhizobial infection and nodule formation (**Figure 3A and Supplementary Figure S3A**). *M. sativa* seedlings were placed onto the Pa-Gr system and inoculated with the engineered *S. meliloti* strains. The system was maintained in a growth chamber to facilitate plant development and symbiotic interaction. After 3-14 dpi, the surface-associated bacteria were removed from the roots using PBS buffer containing 0.02% Silwet. The reporter expressions were then quantified using the FACS platform and observed under a confocal microscopy. Since *S. meliloti* does not naturally form a functional symbiosis with *Lotus japonicus* (Yu and Zhu 2024), their interaction was used as a negative control.

**Figure 3.**
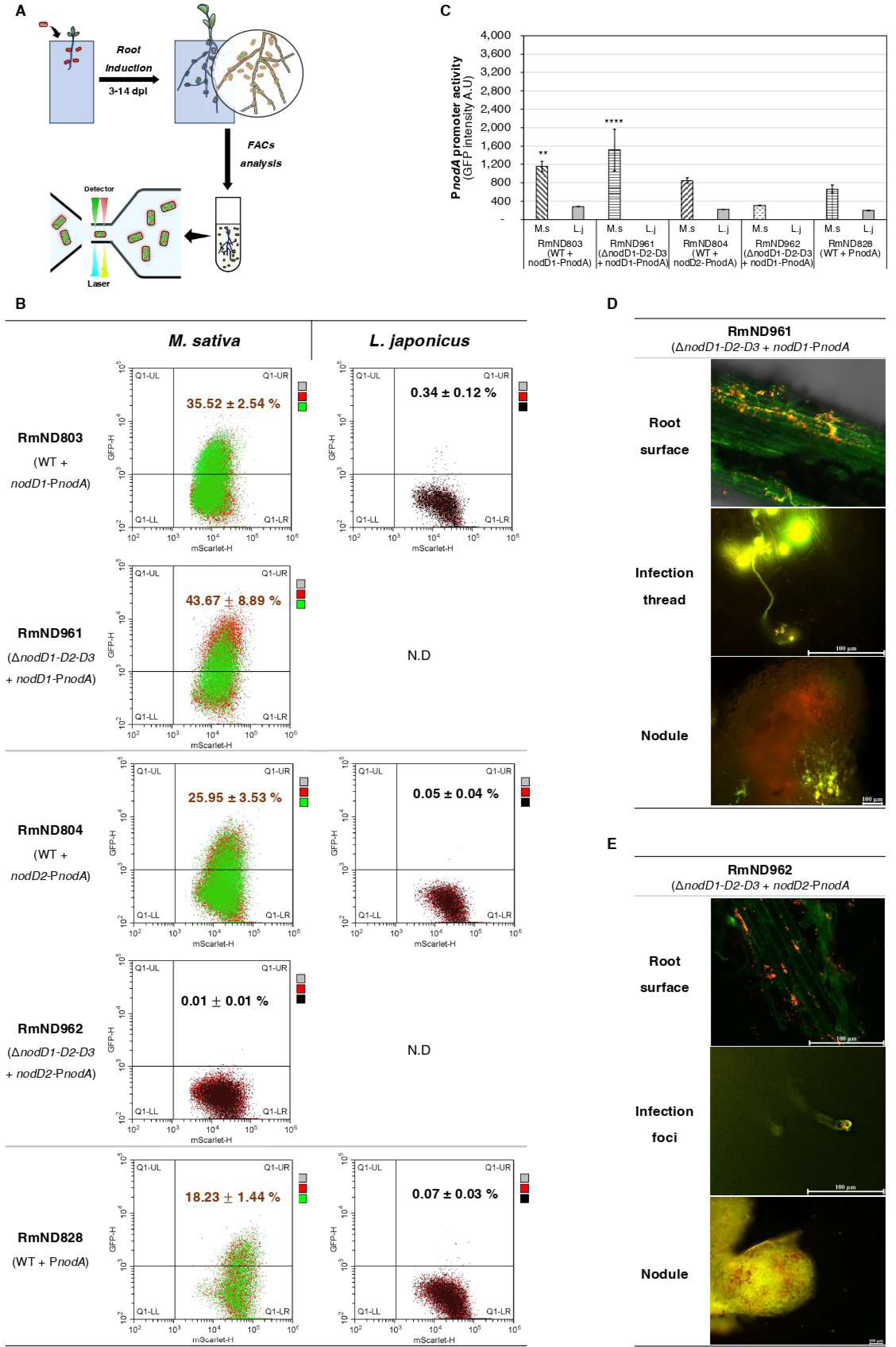
Root induction approach for engineered *S. meliloti* strains harboring the *nod* reporters. (A) Schematic of the paper-based growth (Pa-Gr) system used for root induction, followed by FACs analysis for the biosensor detections. (B) FACs analysis of engineered strains harboring the *S. meliloti nod* reporters collected from roots of *Medicago sativa* and *Lotus japonicus* after 5 days of root induction. The gating strategy established during in vitro experiments was applied to assess biosensor activation. Data represent the proportion of sfGFP-positive cells (mean ± standard deviation) from three independent experiments, each analyzing 10,000 mScarlet-I-expressing bacteria. (C) Quantification of P*nodA* activation in the engineered strains, shown as green fluorescence intensity from FACS analysis. The control strain RmND828 (WT with P*nodA*) was included for comparison. Data represent the green fluorescent intensity (mean ± standard deviation) from three independent experiments. (D-E) Visualization of roots, infection threads and nodules of *M. sativa* infected with RmND961 (triple *nodDs* mutant carrying the *nodD1*-P*nodA*) and RmND962 (triple *nodDs* mutant carrying the *nodD2*-P*nodA*). Symbiotic phenotype characterization of strain RmND961 was performed at 11 dpi and of RmND962 at 28 dpi. *M. sativa* roots, infection threads, and nodules were visualized and analyzed using confocal microscopy. Yellow fluorescence indicates co-expression of both mScarlet-I and sfGFP.

During the root induction, the activation of plant signal biosensor was observed in most engineered strains in the *M. sativa* roots, but not in the *L. japonicus* (**Figure 3B)**. At the 5 days of the root induction, 35.52 ± 2.54 % of RmND803 population (WT with the *nodD1*-P*nodA*) and 43.67 ± 8.89 % of RmND961 population (three *nodDs*-mutant with *nodD1*-P*nodA*) exhibited high sfGFP expression levels, compared to approximately 18.23 ± 1.44 % of the control strain RmND828 (WT with the P*nodA*) (**Figure 3B)**. In contrast, no P*nodA* activity was detected in these strains under the *L. japonicus* root induction. Based on the sfGFP intensity, the P*nodA* activities in RmND803 and RmND961 were approximately 2-fold higher than in RmND828, which lacked the *nodD1* component of the biosensor (**Figure 3C**). These activities showed no significant changes over the 14-day period of root induction (**Supplementary Figure S3B**). Compared to the luteolin-induced expression, both RmND803 and RmND961 showed significantly elevated P*nodA* activity in response to *M. sativa* root exudates **(Figure 3B-C and 1C-D)**. However, these activities remained lower than those observed under the *M. sativa* seedling induction **(Figure 3B-C and 2B-D)**.

Among engineered strains carrying the *nodD2*-driven reporter, only RmND804 (WT with the *nodD2*-driven P*nodA* reporter) exhibited detectable P*nodA* activity during the *M. sativa* root induction (**Figure 3B-C and Supplementary Figure S3B-C**). Approximately 26% of RmND804 population showed high sfGFP expression, compared to 18 % in the control strain RmND828 (P*nodA* promoter alone). The reporter assays indicated that the *PnodA* activity was approximately 1.3-fold that of RmND828 (**Figure 3C**). Compared to trigonelline, *M. sativa* root exudates induced greater P*nodA* activity in RmND804 (**Figure 3C**, **Figure 2D and Figure 1F**). RmND962 (triple *nodDs* mutant with the *nodD2*-P*nodA*) showed insignificant P*nodA* activity, even up to 36 days post-inoculation (**Figure 3B-C and Supplementary Figure S3C**). Altogether, consistent with results observed using seedling induction, these data indicated a modest contribution of *nodD2* to root-based induction of P*nodA* that was also dependent on the presence of other *nodD* proteins in the background genome.

Additionally, the Pa-Gr system enabled the tracking of the microbe-plant interaction. In the symbiosis between *M. sativa* and RmND961 (Δ*nodD1-nodD2-nodD3* mutant with *nodD1*-P*nodA*), we observed P*nodA* expression on the root surface, within the penetrating infection threads (ITs), and in nodule primordia, but not within the mature nodules (**Figure 3D**). In contrast, the betaine biosensor *nodD2*-P*nodA* expression was not observed in the *M. sativa* roots after inoculation with *S. meliloti* RmND962, even up to 36 dpi, despite the strain’s presence on the root surface (**Figure 3E**). Only a few small and non-functional bumps were observed, with no evidence of IT formation or pink, nitrogen-fixing nodules. The results emphasized a minimal role for NodD2 in the invasion of rhizobia, IT formation, and nodule development in the *S. meliloti-M. sativa* symbiosis.

Those achievements suggested that the root induction approach utilizing the Pa-Gr system was effective in enabling *in planta* measurements of the expression of both *nodD1*-P*nodA* and *nodD2*-P*nodA* reporters in engineered strains. Also, this method allowed for real-time observation of the interaction events between the microbes and host plants.

### 3.5 Screening for rhizobia and non-rhizobia with capable of expressing *S. meliloti nod* genes in response to plant signals

The *nodD1*-P*nodA* reporter exhibited robust expression independently, so we wished to explore its efficacy for investigating the capacity to induce NF gene expression across a broad range of microbes. To evaluate its applicability, we introduced the biosensor into several model diazotrophs of interest for their ability to interact with cereal crops (Barnett and Long 2015). These included two cereal-associated rhizobia (*Rhizobium* sp. IRBG74 (Ketelboeter et al., 2023), and *Azorhizobium caulinodans* (Senthilkumar et al. 2008)) and three non-rhizobial diazotrophs (*Klebsiella oxytoca* (El-Khawas & Adachi, 1999; Fujii et al., 1987), *Herbaspirillum seropedicae* (James et al. 1997) and *Azospirillum brasilense* (Schnabel and Sattely 2021)) (**Figure 1B**). As a control, we used the engineered strain *S. meliloti* RmND877 (Geddes et al. 2021), which lacks the 1.35-Mb symbiotic megaplasmid pSymA (including all *nod* genes) but carries the *nodD1*-P*nodA* reporter. These engineered strains were then coated onto *M. sativa* seeds and co-incubated in a plant growth chamber (**Figure 2A**). After 2-d induction, the bacteria were collected, and reporter expression was detected using the FACS platform.

The FACs results indicated the activities of *nodD1*-P*nodA* in various types of microbes after two days of germination induction with *M. sativa* seeds (**Figure 4A**). Among the two model rhizobial strains tested, *Rhizobium* sp. IRBG74 exhibited a sfGFP-positive population of 15.36 ± 2.74%, while the control strain *S. meliloti* RmND877 showed a significantly higher activation level of 38.68 ± 3.44% in response to germinating *M. sativa* seed exudates. In contrast, *A. caulinodans* ORS571 displayed no detectable reporter activity under the same conditions. In the model non-rhizobia, a sfGFP-positive population of *A. brasilense* Sp7 (31.79 ± 4.69 %) was observed after 2-d induction. Meanwhile, no sfGFP expression was detected in *K. oxytoca* M5A1 and *H. seropedicae* LR15 populations. The biosensor quantification assays confirmed the activation of the flavonoid-dependent biosensor in both *Rhizobium* sp. IRBGG74 and *A. brasilense* Sp7 (**Figure 4B**). The P*nodA* activities of these strains under the germinating *M. sativa* seed induction were approximately 3.8-fold and 1.4-fold lower, respectively, than that of the control strain RmND877. The achievement suggested that the flavonoid reporter *nodD1*-P*nodA* could be used as a selectable marker for high-throughput screening engineered N_2_-fixing bacteria with the capacity to utilize NF signaling to communicate with plants.

**Figure 4.**
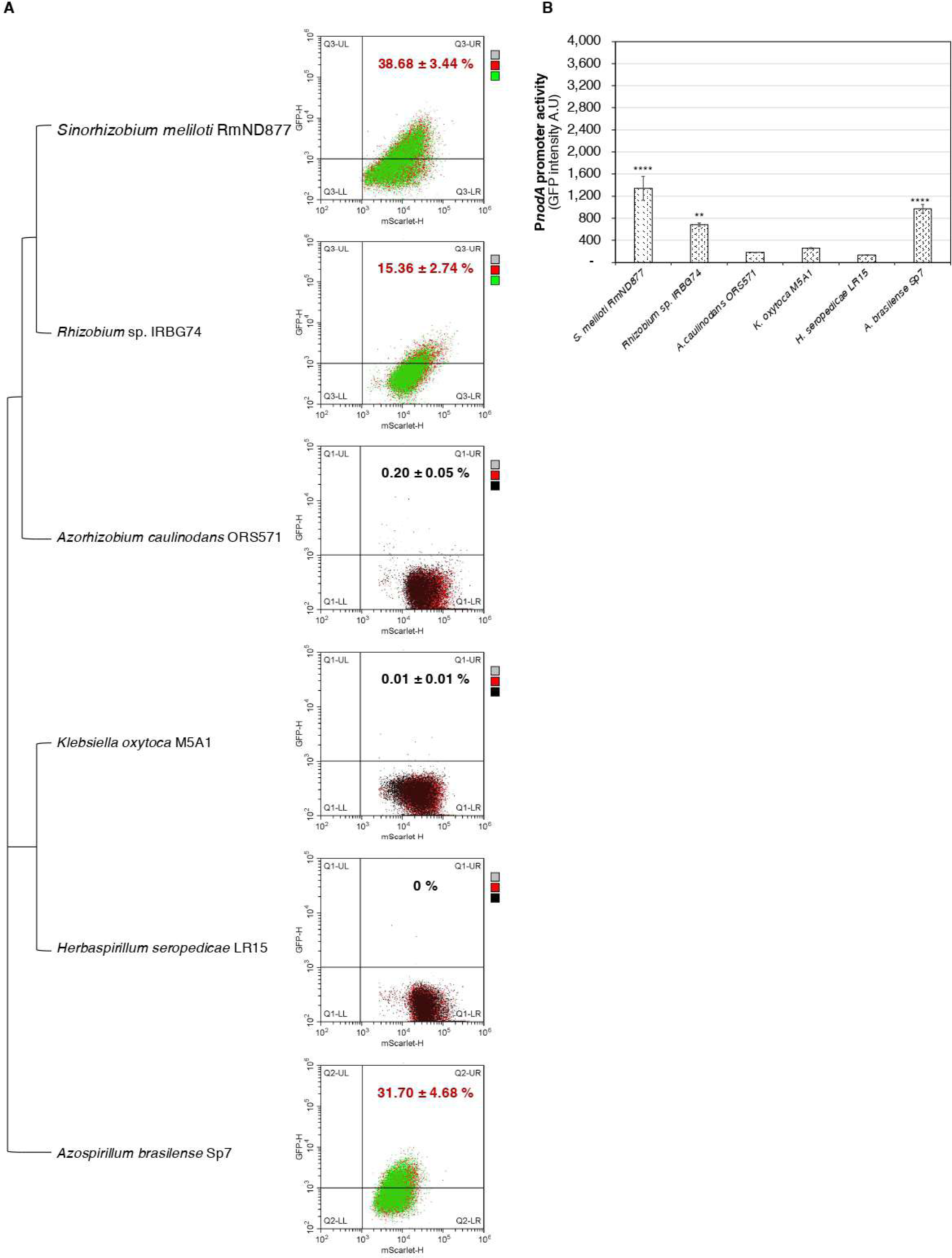
Functional expression of *nodD1*-P*nodA* in rhizobia and non-rhizobia. (A) 16S rRNA Phylogenetic tree and FACs analysis of the flavonoid biosensor expression in various microbes after 2 days of germinating *M. sativa* seed induction. The phylogenetic tree includes *Sinorhizobium meliloti* RmND877 (positive control strain), *Rhizobium* sp. IRBG74, *Azorhizobium caulinodans*, *Klebsiella oxytoca*, *Herbaspirillum seropedicae*, and *Azospirillum brasilense* harboring the *nodD1*-P*nodA* reporter. The same gating strategy was applied to identify sfGFP-positive cells. Data represent the mean ± standard deviation of sfGFP-positive populations from three independent experiments, each analyzing 10,000 mScarlet-I-expressing cells. (B) Fluorescence readout for activation of *nodD1*-*PnodA* reporter based on the FACs analysis. Non-sfGFP-expressing strains were served as negative controls. Data represent the green fluorescent intensity (mean ± standard deviation) from three independent experiments.

## 4 DISCUSSION

Engineering synthetic nitrogen (N_2_)-fixing symbioses in cereals requires precise tools to identify compatible and engineerable microbial partners (Peck, Fisher, and Long 2006). In this study, we developed and validated two plant signal biosensors, including a flavonoid-responsive *nodD1*-P*nodA* and a betaine-responsive *nodD2*-P*nodA* (**Figure 1A**), using the *Sinorhizobium meliloti*-*Medicago sativa* symbiosis model system. The *nodD1*-P*nodA* reporter system exhibits robust and independent activation in response to plant-derived flavonoid signals. In contrast, *nodD2*-P*nodA* showed only minimally detectable activity under similar conditions. These biosensors enabled functional screening of rhizobial and non-rhizobial strains for their ability to perceive and respond to host-derived signals by producing *S. meliloti* NFs. The system provides a platform for identifying microbes suitable for synthetic symbiosis engineering and offers insights into the distinct regulatory roles of NodD1 and NodD2 in *S. meliloti nod* gene activation in the early stages of *S. meliloti-M. sativa* interaction.

Under *in vivo* induction, *S. meliloti* strains carrying inducible biosensors responded specifically to individual purified plant-derived signals (**Figure 1C-F**). In *S. meliloti*, NodD1 was activated by flavonoids such as luteolin and methoxychalcone, whereas NodD2 responds to plant betaines like trigonelline and stachydrine, as well as the flavonoid 4,4’-dihydroxy-2’-methoxychalcone (Barnett and Long 2015; Maxwell et al. 1989; Phillips et al. 1992). After 4-h induction with 10 µM luteolin, both RmND803 (WT with *nodD1*-P*nodA*) and RmND961 (Δ*nodD1-D2-D3* mutant with *nodD1*-P*nodA*) exhibited strong flavonoid-dependent NodD1-mediated activation (**Figure 1C-D**), consistent with the luteolin-enhanced promoter binding study (Peck et al. 2013). Similarly, 10 mM trigonelline induced detectable activation of the *nodD2*-P*nodA* reporter in RmND804 (WT background) (**Figure 1E-F**), confirming NodD2-mediated activation, though to a much lesser magnitude. These results validated the functionality of our biosensor system in response to individual plant signals.

To validate the biosensor’s functionality in a physiologically relevant context, we employed germinating seed and root induction approaches for the first time (**Figure 2A and 3A**). These approaches enabled dynamic assessment of biosensor activation in response to natural plant-derived signals during the early stages of *S. meliloti-M. sativa* symbiosis. Notably, flavonoid-dependent *nodD1*-P*nodA* activity was observed in response to germinating *M. sativa* seed exudates and root exudates in both WT and Δ*nodD1-D2-D3* mutant backgrounds (**Figure 2 B-D and 3 B-C**). The activity was highest in response to germinating seed exudates (**Figure 2D**), followed by root exudates (**Figure 3C**), and lowest with luteolin induction (**Figure 1D**). This enhanced activation may be attributed to the presence of a diverse cocktail of nodulation-inducing molecules that vary across different developmental stages of the host plant (Liu and Murray 2016). Metabolomic analyses of germinating *M. sativa* seed exudates have identified hyperoside, luteolin-7-glucoside, luteolin, and chrysoeriol as the four most abundant flavonoids (Hartwig et al. 1990; Compton et al. 2020). These compounds are known to play critical roles in early symbiotic signaling and microbial interactions in the rhizosphere. At the later growth stages, recent flavonoid metabolome profiling of *M. truncatula* roots identified a total of 335 flavonoid compounds with 98 differentially accumulated flavonoids in response to *S. meliloti* inoculation (Shen et al. 2025). Our findings highlighted the primary role of the flavonoid-responsive regulator NodD1 in activating *S. meliloti nod* gene expression and NF biosynthesis during the early stages of *S. meliloti-M. sativa* symbiosis, even in the absence of NodD2 and NodD3 (**Supplementary Figure S2 and Supplementary Figure S3B**). The independent expression of the flavonoid biosensor facilitated its applications in studying *S. meliloti* nodulation gene expressions in various rhizobia and non-rhizobia and establishing host specificity between these microbes and target crops.

Regarding the functionality of the betaine-responsive biosensor *nodD2*-P*nodA*, it exhibited its highest level of activation in the presence of *nodD1-D3* in wild-type *S. meliloti* (**Figure 2C, 3B, 3C and Supplementary Figure S3B**). In the presence of wild-type *nodD*s, the *nodD2-*P*nodA* sensor still showed greater induction than the P*nodA* sensor alone. A plausible explanation for this observation is the presence of 4,4’-dihydroxy-2’-methoxychalcone (DHMC) in the seedling and root exudates of *M. sativa* (Maxwell et al. 1989). Notably, even at a low concentration of approximately 2 nM, DHMC was sufficient to trigger NodD2-mediated transcriptional activation (Phillips et al. 1992). Taken together, these data indicate that effective *nodD2*-mediated induction of *PnodA* is at least partially dependent on either *nodD1* or *nodD2.* Other studies have also reported NodD2 functions more effectively as an activator when NodD1 or NodD3 is present, highlighting its synergistic regulatory capacity (Honma, Asomaning, and Ausubel 1990). Despite its limitations, it still offers potential as a tool for customized microbial engineering and environmental sensing. Betaine-related compounds, such as trigonelline and stachydrine, not only act as *nod* gene inducers in *S. meliloti* (Phillips et al. 1992) but also serve as stress protectants for bacteria and plants against salinity, drought, flooding, heavy metals, cold, and heat stresses (Hernandez-Leon and Valenzuela-Soto 2023; Alloing et al. 2006; Boscari et al. 2006). These make betaine-responsive biosensors attractive tools for designing stress-resilient rhizobial strains.

Using symbiotic signal biosensors, we elucidated the distinct regulatory roles of NodD1 and NodD2 in NF production during early stages of the *S. meliloti-M. sativa* symbiosis. NodD1 emerged as the principal activator of *S. meliloti nod* gene expression, functioning effectively in the rhizosphere, within infection threads, and in in nodule primordia, but showing limited activity in mature nodules (**Figure 3D**). In contrast, NodD2 showed limited transcriptional activation capacity and was unable to initiate effective symbiotic signaling on its own. Our findings indicated that inoculating *S. meliloti* with NodD2 alone leads to delayed nodulation and formation of white, non-functional nodules in *M. sativa* (**Figure 3E**). Consistent with these observations, a previous study demonstrated that mutations in NodD1 severely disrupt nodulation on *M. sativa*, whereas NodD2 mutations had minor effects (Honma and Ausubel 1987). NodD2-mediated activation is limited in *M. sativa* due to insufficient levels of NodD2-specific inducers in root exudates. In interactions with *Melilotus alba*, NodD2-mediated activation was detectable in the presence of root exudates and the Δ*nodD1-D3* mutant was still able to nodulate (Honma, Asomaning, and Ausubel 1990), suggesting a conditional role for NodD2. Beyond *S. meliloti*, NodD2 exhibits contrasting roles in other rhizobia. In *Mesorhizobium loti (Mt)–Lotus japonicus* symbiosis, for example, NodD2*_Mt_* was found to play a major role in *M. loti nod* gene induction within the rhizosphere and developing nodules, while NodD1*_Mt_* is primarily active in root hair infection threads (Kelly et al. 2018). In *Rhizobium tropici* (*Rt*), NodD2*_Rt_* can even activate *R. tropici nod* genes under abiotic stress in a flavonoid-independent manner, expanding host range and enhancing stress resilience (Del Cerro et al. 2017). Overall, these findings indicate that NodD2 acts as a conditional activator rather than a primary one, regulating *nod* gene expression in response to host-specific and environmental conditions.

Finally, as proof of concept, we utilized the flavonoid biosensor *nodD1*-P*nodA* for identifying model rhizobia and non-rhizobia that can produce NF signals in response to flavonoid signals (**Figure 4**). Among the tested rhizobia and non-rhizobia, *Rhizobium* sp. IRBG74 and *Azospirillum brasilense* Sp7 exhibited significant reporter activation. In comparison to the control strain *S. meliloti* RmND877, the reporter activities in *Rhizobium* sp. IRBG74 and *Azospirillum brasilense* Sp7 were 3.8-fold and 1.4-fold lower, respectively (**Figure 4B**). The differential activations might reflect variations in flavonoid uptake efficiency and regulatory mechanisms controlling the *nod* gene expression in these backgrounds. For instance, *S. meliloti* could absorb luteolin at rates nearly twice that of *Rhizobium leguminosarum* biovar *viciae* and up to 10-fold greater than non-rhizobia like *Agrobacterium tumefaciens, A. rhizogenes, Pseudomonas syringae, E. coli* (Hubac et al. 1993). The notable reporter activation in *A. brasilense* background could be related to pre-existing ecological interactions with flavonoids, for instance, enhanced secretion of flavonoids by legumes (Dardanelli et al., 2008). Both *Rhizobium* sp. IRBG74 and *A. brasilense* are endophytic bacteria associated with cereal crops, known to enhance plant growth and yield (Igiehon and Babalola 2018; Ryu et al. 2020). Although the reporter activity in these endophytes was lower than that of the native *S. meliloti* strains, the biosensor’s sensitivity was sufficient to detect potential N_2_-fixing diazotrophs. To further quantify microbial competition, fitness, and colonization dynamics within diazotrophic communities in cereals, the Plasmid-ID system could be integrated into the plant signal-dependent biosensor systems (Schumacher et al. 2025). Moreover, the compatibility of the flavonoid biosensor with high-throughput screening technologies, such as VECTOR (Versatile Engineering and Characterization of Transferable Origins and Resistance) (Williamson et al. 2025), fluorescence-activated cell sorting (FACS) (Jorrin et al. 2024), and fluorescence-activated droplet sorting (FADS) (Luu et al. 2022), offers a powerful approach for isolating and selecting high-performing strains within microbial consortia or synthetic communities.

Together with the generation of nodule-like structures with NF receptors in cereal crops (Guo et al. 2023), the symbiotic signal biosensors represent a promising tool for enabling both symbiotic and associative nitrogen fixation in cereal crops, contributing to the broader goal of sustainable agriculture. We demonstrated that *S. meliloti* NodD1 is functional in important cereal-colonizing diazotrophs, including *Rhizobium* sp. IRBG74 and *A. brasilense* Sp7 and can coordinate host-specific induction of gene expression in response to *M. sativa*. This strategy could be extended to cereals by designing biosensors that respond to cereal-specific root exudates. Alternatively, NodD3 can be used to drive constitutive NFs induction to “short circuit” the first steps of signaling (Haskett et al. 2025). Several studies have demonstrated that engineered barley can excrete rhizopine, a plant-derived signaling molecule, to activate gene expression in bacterial colonies colonizing barley roots (Haskett et al., 2025; Haskett et al., 2022; Ryu et al., 2020), which could provide an alternative inducing system for NF synthesis either directly through refactoring *nod* gene regulation or indirectly through controlling *nodD*s. In any case, major efforts have been placed in cereal crops to engineer the perception of symbiotic signals and the concomitant signal transduction through the “common symbiotic pathway” in the past decade (Rübsam et al. 2023; Jhu and Oldroyd 2023). Since rhizobia evolved for millions of years with their host-legumes to facilitate accommodation by plant immunity within their intimate symbiotic relationship (Wang et al. 2020), we anticipate a need to address the engineering of suitable symbionts for synthetic cereal root nodules, from chassis with pre-existing ecological interactions (Rübsam et al. 2023; Jhu and Oldroyd 2023). This work contributes to this challenge by developing tools and new insights for transferring the first steps in symbiotic signaling to cereal-colonizing non-rhizobia diazotrophs.

## 5 CONCLUSIONS

In this study, we established a biosensor-based platform for detecting plant-derived signals and evaluating microbial compatibility in the context of engineering synthetic nitrogen-fixing symbioses. By constructing flavonoid- and betaine-responsive biosensors (*nodD1*-P*nodA* and *nodD2*-P*nodA*), we enabled precise monitoring of *Sinorhizobium meliloti nod* gene activation in the native and other microbial candidates. These biosensors demonstrated specificity and functionality under both *in vitro* and *in planta* conditions. As a proof of concept, the flavonoid biosensor *nodD1*-P*nodA* was successfully applied to identify rhizobial and non-rhizobial strains capable of producing *S. meliloti* NFs, including *Rhizobium* sp. and *Azospirillum brasilense* backgrounds. The integration of our symbiotic signal biosensors with high-throughput screening technologies will provide a scalable strategy for selecting high-performing microbial strains from large consortia. This approach establishes a foundation for accelerating the development of synthetic symbioses in cereal.

## Acknowledgements

This work was supported by a New Innovator in Food & Agricultural Research (FFAR) grant to BAG ID: FF-NIA21-0000000061 and the Richard and Linda Offerdahl Faculty Fellowship to Barney A. Geddes. We thank Melanie Barnett and Sharon Long for sharing strain resources. We are grateful to Scott Hoselton and Kaycie Schmidt at the Dr. Thomas Glass Biotech Innovation Core, in the Department of Microbiological Sciences, North Dakota State University (NDSU) for technical support with flow cytometry. We thank Dr. Pawel Borowicz from Department of Animal Sciences, NDSU, for assistance with the Mica Microhub system. We are also grateful to Megan Ramsett in the Department of Microbiological Sciences, NDSU for administrative and technical contributions to this work. Finally, we thank Maryam Khan in the same department for experimental support in setting up the paper-growth system.

## Author Contributions

*Chinh. X Luu*: conceptualization, methodology, formal analysis, investigation, data curation, writing - original draft. *Barney A. Geddes:* conceptualization, validation, supervision, resources, writing - review and editing, project administration.

## Data Availability Statement

The data supporting the findings of this study are included in the supplementary information files.

## Abbreviations

NFs: Nod factors
*nod*: gene encoding NFs
sfGFP: superfolder green fluorescent protein
FACs: fluorescence-activated cell sorting
*sfgfp*: gene encoding superfolder green fluorescent protein
WT: wild-type

